# CRISPR-Cas9 mediated deletions of *FvMYB46* reduces fruit set and biosynthesis of flavonoids in *Fragaria vesca*

**DOI:** 10.1101/2024.08.07.607017

**Authors:** Arti Rai, Magne Nordang Skårn, Abdelhameed Elameen, Torstein Tengs, Mathias Rudolf Amundsen, Oskar S. Bjorå, Lisa K. Haugland, Igor A. Yakovlev, May Bente Brurberg, Tage Thorstensen

**Author notes:** **Correspondence:** Tage Thorstensen.

## Abstract

Secondary metabolites produced by the phenylpropanoid pathway, which is regulated by transcription factors of the MYB family, play crucial roles in this early phase of fruit development. The MYB46 transcription factor is a key regulator of secondary cell wall structure and lignin and flavonoid biosynthesis in many plants, but little is known about its activity in flowers and berries in *F. vesca*. For functional analysis of FvMYB46, we designed a CRISPR-Cas9 construct with an endogenous *F. vesca* specific U6-promoter for efficient and specific expression of two gRNAs targeting the first exon of *FvMYB46*. This generated mutants with an in frame 81-bp deletion of the first conserved MYB-domain or an out of frame 82-bp deletion potentially knocking out the gene function. In both types of mutant plants, pollen germination and the frequency of flowers developing to mature berries was significantly reduced compared to wild type. Transcriptomic analysis of flowers demonstrated that FvMYB46 is positively regulating the expression of genes involved in pollen germination, homeostasis of reactive oxygen species (ROS) and the phenylpropanoid pathway, including secondary cell wall biosynthesis and flavonoid biosynthesis, while has a negative impact on carbohydrate metabolism. In FvMYB46-mutant flowers, the flavonols and flavan-3-olscontent, especially epicatechin, quercetin-glucoside and kaempferol-3-coumaroylhexoside were reduced, and we observed a local reduction of lignin content in anthers. Together these results suggest that MYB46 control fertility and efficient fruit set by regulating cell wall structure, flavonoid biosynthesis, carbohydrate metabolism and ROS-signaling in flowers and early fruit development in *F. vesca*.

## Introduction

Successful fertilization and fruit set, where the ovary develops to a young fruit, are important for crop yield as they determine the number and size of fruits and seeds in plants (Ruan et al., 2012). Development and differentiation of anthers and pistils, the male and female reproductive organs respectively, is precisely regulated for fertilization to succeed. In cultivated strawberry (*Fragaria × ananassa*) which is a commercially important soft fruit crop producing berries rich in beneficial vitamins, nutrients and phenolic compounds, both environmental and genetic factors influence anther development and pollen quality important for high fruit setting rate (Kronenberg, 1959). The diploid woodland strawberry *Fragaria vesca* is used as a model plant for gene functional studies and flower and fruit development because it has a small genome, is easily transformed and contains few duplicated regions compared to the genomically more complex octoploid cultivated strawberry (Shulaev et al., 2011). Several transcriptomic analyses of *F. vesca* have given molecular insight into fertilization and the early-stage fruit development (Hollender et al., 2014; Kang et al., 2013; Shahan et al., 2018). Recently, gene editing using CRISPR-Cas9 has successfully been used for functional studies of some of the genes involved in fruit development in both *F. vesca* and *F. × ananassa* (L. Guo et al., 2022; López-Casado et al., 2023; Mao et al., 2022; Martín-Pizarro et al., 2019).

The phenylpropanoid pathway produces a diverse group of secondary metabolites including precursor compounds for flavonoid and lignin biosynthesis. The precursors are then modified by enzymes in the flavonoid or lignin biosynthetic pathways to produce metabolites involved in protection against biotic and abiotic stress, plant reproduction and cell wall structure (Anwar et al., 2021; Yadav et al., 2020). Secondary cell walls (SCW) consist of cellulose, hemicellulose and lignin and forms between the primary cell wall and plasma membrane and is crucial for mechanical strength and maintaining cell shape and function in specialized cells such as fibers and xylem.deposition of lignin and formation of secondary cell walls also improve water conductivity and provides stress tolerance in plants. In anthers, formation of lignified, cellulosic secondary wall thickenings in cells of the endothecium is required for anther dehiscence, creating a pressure for pollen exposure and dispersal. This process is precisely controlled and disruption in development of these thickenings prevents anther dehiscence and causes male sterility (Dawson et al., 1999; Keijzer, 1987). The receptacle, which is a modified stem tip in strawberry, is topped with dozens of pistils, each containing an individual carpel with ovary and ovule. Successfully fertilized ovules develops seeds and produces the phytohormones auxin and gibberellins which stimulates the receptacle to enlarge into the fleshy structure known as the berry (Hollender et al., 2012; Kang et al., 2013; Z. Liu et al., 2020; Tian et al., 2022). Auxin also signals the ovary wall (carpel wall) to enlarge and develop into dry achenes, which is probably due to cell wall synthesis and extensive lignification (Zhou et al., 2023). This positive control of early fruit development by auxin ensures that fruit set only occurs after successful fertilization.

MYB transcription factors bind to specific DNA sequences in the promoter regions of target genes that influence numerous processes, including secondary cell wall biosynthesis, abiotic stress tolerance, resistance to biotrophic and necrotrophic pathogens and flower organ development (Wang et al., 2021; Xiao et al., 2021). Several studies have shown that MYB-transcription factors regulate phenylpropanoid metabolism important for fertilization and early fruit development (J. Liu et al., 2015; Pratyusha & Sarada, 2022). The R2R3-type MYB26 transcription factor regulates anther dehiscence important for successful fertilization by inducing secondary thickening through the NAC transcription factors NST1 and NST2 in *Arabidopsis* (Mitsuda et al., 2005; Steiner-Lange et al., 2003; Yang et al., 2017). NST1 and NST2 have together with NST3/SND1 (SECONDARY WALL-ASSOCIATED NAC DOMAIN PROTEIN1 (SND1), also been shown to be involved in secondary cell wall synthesis in fibers and xylem of inflorescence stems (Zhong & Ye, 2015).

The MYB46-transcription factor is a key player in the transcriptional network regulating secondary wall biosynthesis in xylem cells of inflorescence stems in *Arabidopsis* where it acts redundantly with MYB83 (McCarthy et al., 2009; Zhong et al., 2007). Both MB46 and MYB83 are directly regulated by several NAC TFs, such as VND6 and VND7, NST3/ANAC012/SND1, NST1 and NST2 (Ko et al., 2014; Zhong et al., 2007). MYB46 directly binds and activates transcription factor genes and genes involved in biosynthesis of cellulose, hemicellulose and lignin (Kim et al., 2013, 2014; Ko et al., 2014). The role of MYB46 in secondary cell wall polymerization and biosynthesis of cellulose and lignin has been demonstrated for several species, including birch and apple, where MYB46 activity also improves salt and osmotic stress tolerance (Chen et al., 2019; H. Guo et al., 2017). In *F. vesca*, FvMYB46 was recently found to be a direct target of the NAC transcription factor FvVND4c and to regulate SCW thickening and flavonoid accumulation when overexpressed (Zhang et al., 2023).

In this study we show that CRISPR-Cas9-mediated in-frame 81-bp- or out-of-frame 82-bp deletions of the first exon of *FvMYB46*, both generate plants with reduced pollen germination and frequency of flowers developing to mature berries. Transcriptome analysis by RNA sequencing demonstrated that *FvMYB46* positively regulates the expression of genes involved in the phenylpropanoid pathway, secondary cell wall biosynthesis, stress tolerance and ROS-signaling in flowers. We also observed an upregulation of genes involved in carbohydrate metabolism and signaling. Metabolite profiling and histological stainings confirmed that FvMYB46 is an activator of flavonoid biosynthesis in flowers and lignin content in anthers. Our work demonstrates that FvMYB46 control efficient fertilization and fruit set by biosynthesis of flavonoids and cell wall components and regulating genes involved in ROS signaling and carbohydrate metabolism in *F. vesca*.

## Results

### Expression of FvMYB46 in different F. vesca tissues

To identify *MYB46* homologues in *F. vesca*, we used the *MYB46*-nucleotide sequence from *Arabidopsis* (AT5G12870) for blastn searches against the *F. vesca* nr genome sequence database at NCBI. This identified the two *F. vesca* transcripts FvH4_3g28890 and FvH4_7g01020 as potential *AtMYB46* homologs. Multiple sequence alignment of the protein sequences of the potential *F. vesca* MYB46 homologs with MYB46 and MYB83 transcription factors (Figure 1 a) with known function from other plants showed that FvH4_3g28890 clustered with the MYB46 group and was named FvMYB46, while FvH4_7g01020 clustered with the MYB83 group and named FvMYB83 (Figure 1 b). The expression profile of *FvMYB46* in different *F. vesca* tissues obtained by qRT-PCR analysis, showed a very low expression level of *FvMYB46* in roots, young leaves and mature berries, and slightly higher in mature leaves and seedlings. Expression was moderate in open flowers, and highest in inflorescence stems and young berries (Figure 1 c). Querying the eFP (electronic Fluorescent Pictograph) browser, showed that *FvMYB46* has the highest expression in anthers at stage 11 of flower development before fertilization, and in the carpel walls (ovary walls) in the first stages of fruit development after fertilization (Figure 1 d) (Hollender et al., 2012; Kang et al., 2013).

**Figure 1.**
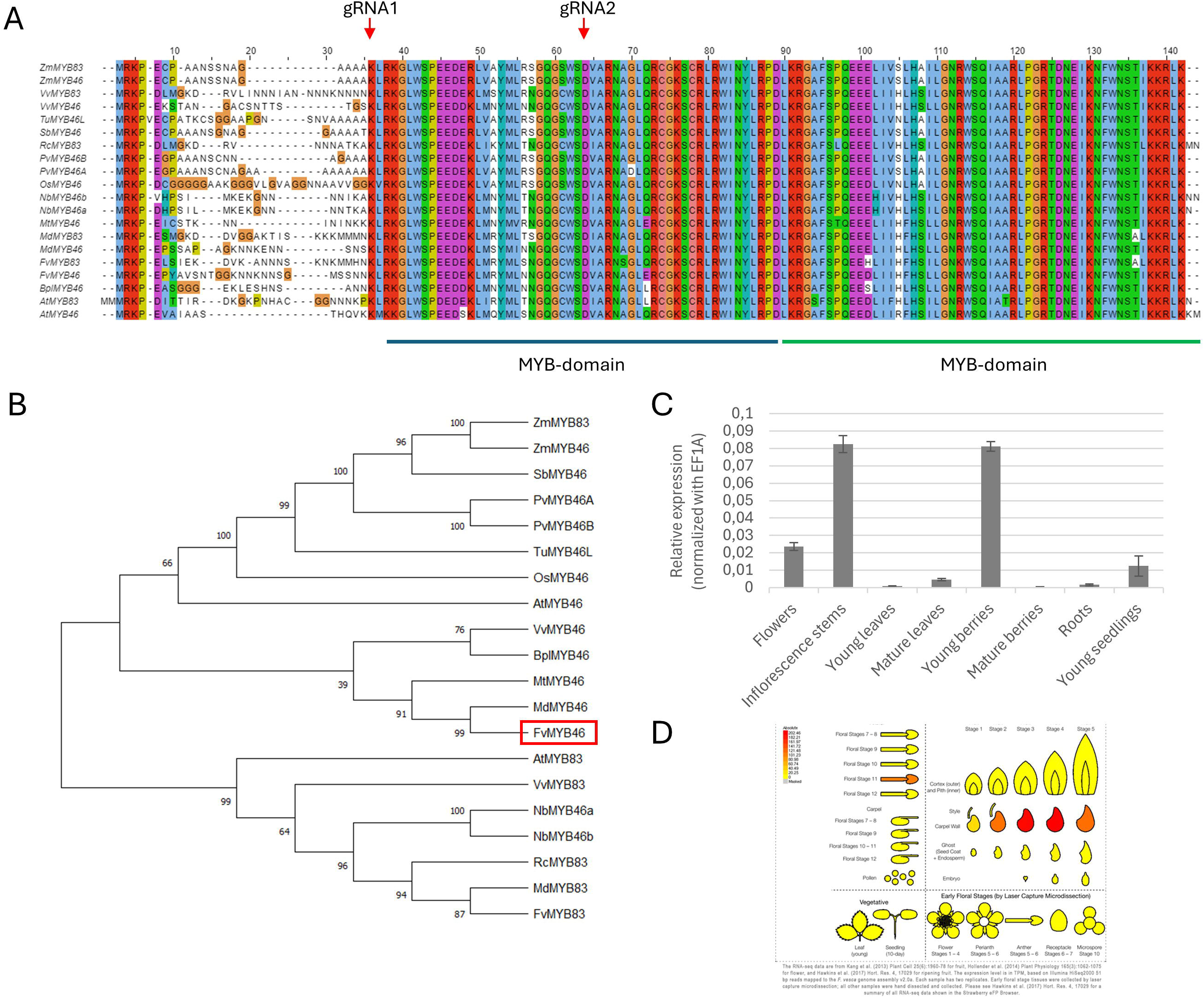
MYB46-homologues from different plant species. A) Alignment of sequences of the N-terminal region comprising MYB-domain 1 and MYB-domain 2 from functionally analyzed MYB46 proteins from different plant species. Arrowheads indicate deletions mediated by gRNA1 and gRNA2. B) Phylogenetic analysis of protein sequences from A) using the neighbour joining method with 1000 bootstrap sampling. C. qRT-PCR of FvMYB46 on cDNA from RNA isolated from different tissues of wild type F. vesca. Expression was normalized with EF1A. D) Expression analysis of FvMYB46 in different organs and stages of flower and fruit development using eFP-browser. Error bars are ± standard error based on 3 biological replicates.

### CRISPR-Cas9 knock out of FvMYB46

We used CRISPR-Cas9 to specifically introduce gene deletions in *FvMYB46* for functional analyses. The AtU6 promoter is frequently used to drive the sgRNA expression for CRISPR genome editing in dicot plants, but endogenous promoters have been shown to be more efficient in many species (Ren et al., 2021; Riu et al., 2023). To identify endogenous FvU6 promoters, we used U6 snRNA genes from *Arabidopsis* and *T. aestivum* in BLAST searches against the *F. vesca* genome. In transient expression analyses the FvU6-1 promoter showed stronger and more robust expression of gRNAs than the AtU6-promoter, hence FvU6-1 was applied for expression of two gRNAs targeting the first exon of *FvMYB46* in CRISPR-Cas9 constructs used for *Agrobacterium* mediated transformation of *F. vesca* (Supplementary material, Figure S1). PCR screening using primers flanking the gRNA target region and direct sequencing of 50 primary transformants (T0) identified three different genotypes containing homozygous deletions of 81 and 82 bp, or bi-allelic 81/82 bp deletions in both the *FvMYB46* gene and cDNA from flowers (Figure 2 a, b and Figure S2 b, c). The wild type full length *FvMYB46* gene encodes a protein of 335 aa, while the 81 bp deletion creates an in-frame deletion of 27 aa, resulting in a putative protein of 308 aa. The 27 aa deletion removes most of the first of two N-terminal MYB/Sant/HTH domains important for DNA-binding and protein-protein interactions. In contrast to the 81bp deletion, the 82 bp deletion creates a putative out-of-frame translation product of 50 aa, where only the first 28 aa are identical to the wild type protein (Figure S2 a), thus containing none of the conserved MYB domains. To avoid any influence on downstream molecular and phenotypic analyses, we removed the T-DNA containing the CRISPR-Cas9-cassette by selfing primary transformants of the T0 generation. Homozygous T1 progeny of the 82-bp out-of-frame deletion (*FvMYB46-82*) and bi-allelic progeny with an in-frame 81bp deletion (*FvMYB46-81*) and an out-of-frame 82-bp (*FvMYB46-81*) of the *FvMYB46* gene, and negative for T-DNA containing Cas9 were used for further studies (Supplementary Figure S2 c).

**Figure 2.**
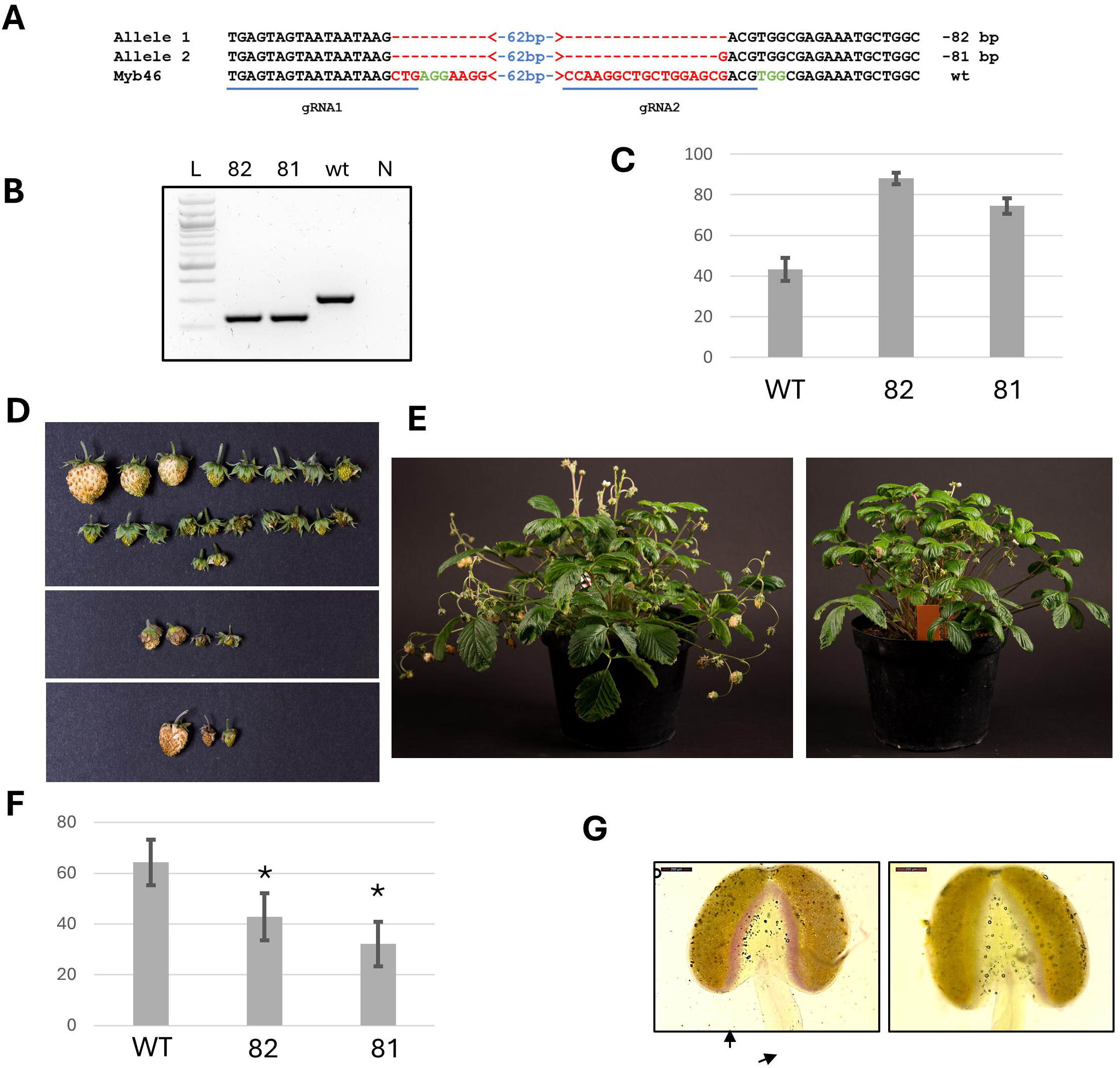
CRISPR Cas9-deletions of FvMYB46 causes reduced fruit set, pollen germination and anther lignification. A) Sequence alignment of the FvMB46-81 and FvMYB46-82 deletions with wt. The two gRNA sequences are underlined. The PAM sequences are colored in green, and deleted region in red. B) Expression of FvMB46 in flowers of WT and mutants, analyzed by RT-PCR using gene specific primers flanking the gRNA1 and gRNA2 target sequences. L, 100 bp DNA ladder; 82, FvMYB46-82 mutant; 81, FvMYB46-81/82 mutant; wt, wild type; N, dH2O neg control. C) Number of mature berries from wild type, FvMYB46-82 and FvMYB46-81/82 mutants were divided by the total number of flowers to determine the fruit set. Graph shows average percent fruit set per plant ± Standard error, n=12. D) Representative picture of fruit yield from wt (top), FvMB46-81/82 (middle) and FvMYB46-82 (bottom) plants. E) Representative picture of wild type (left) and FvMYB-82 (right) plants. F) Frequency of flowers with germinating pollens from wild type and mutants ± Standard error, n=28. Asterisk indicate significant difference from wt based on Anova test (P<0,05). G) Anthers from wild type (left) and FvMYB46-82 (right) plants. Arrows shows stained vascular bundles and endothecium facing the connective tissue.

The specificity of the gRNA-directed deletions was determined by PCR-amplification of the 3 top ranking potential off-target sites for gRNA1 and 1 site for gRNA2, and sequencing of these PCR products. No mutations except for the intended deletion at *MYB46* were found. We also analyzed the RNAseq data for potential mutations in any other sites in the genome but did not identify any off-target mutations or other insertion sites for T-DNA.

### Analyzing fruit set in myb46-plants

The high expression of *FvMYB46* in inflorescence stems, flowers and young unripe berries compared to other organs and development stages, and the specific expression in anthers and ovary walls, suggests a role for *FvMYB46* in fertility or early fruit development. By calculating the frequency of flowers that developed into mature fruits, we observed that the fruit set was significantly lower in the homozygous *FvMYB46-82* and biallelic *FvMYB46-81/82* plants than in the wild type (Figure 2 c). Compared to the wild type, a larger number of the flowers in both mutants were seemingly arrested in the early receptacle fruit development, after which they died and dried up in contrast to flowers that developed into normal fleshy fruits (Figure 2 d, e). Reduced fruit set can be a consequence of reduced pollination caused by reduced pollen viability, pollen tube elongation and fertilization. We observed a reduced number of flowers with germinating pollen in both mutant genotypes compared to the wild type (Figure 2 f), suggesting that FvMYB46 is involved in pollen development and/or viability. Except for lower fruit yield and higher frequency of berries arrested in early fruit development, we did not observe other morphological deviations in *FvMYB46-82* and *FvMYB46-81/82* plants during vegetative growth (Figure 2 e).

### Microscopy analyses of secondary cell walls in inflorescence and anthers

Microscopy analyses of *FvMYB46-82, FvMYB46-81/82* and wild type anthers stained with phloroglucinol-HCl were carried out to study lignin content in the secondary cell walls.

Phloroglucinol-HCl staining of buds before anthesis and open flowers, show modest but specific reduced staining of mutant anthers compared to wild type, especially in endothecium cells facing the connective tissue, and vascular tissue of the filament (Figure 2 g). However, the total lignin content was not significantly different in flowers of wild type and mutants (Figure S3), suggesting that the observed modest reduction in lignin content in the cell walls of anthers is not affected on a global level.

### Transcriptional profiling of FvMYB46-81/82 and FvMYB46-82 flowers

RNA-seq was then performed on open flowers and unripe berries to identify the genetic pathways affected by *FvMYB46* in early fruit development. Comparing differentially expressed genes (DEGs) in *FvMYB46-82* and *FvMYB46-81/82* plants with wild type (FDR <0.01) (Supplementary table 1), we found that 631 of the annotated transcripts mapped to the *F. vesca* v4.0.a2 transcriptome were downregulated and 544 were upregulated in *FvMYB46-82* flowers, while 1247 were downregulated and 925 were upregulated in *FvMYB46-81/82* flowers. 768 genes differently expressed in *FvMYB46-82* flowers were also differently expressed in the FvMYB-81/82 flowers (Figure 3 a). Principal component analysis (PCA) separated wild type and from both the *FvMYB46-82* and *FvMYB46-81/82* mutants which clustered together but formed separate subgroups, suggesting that the gene expression was affected by the mutations (Figure 3 b).

**Figure 3.**
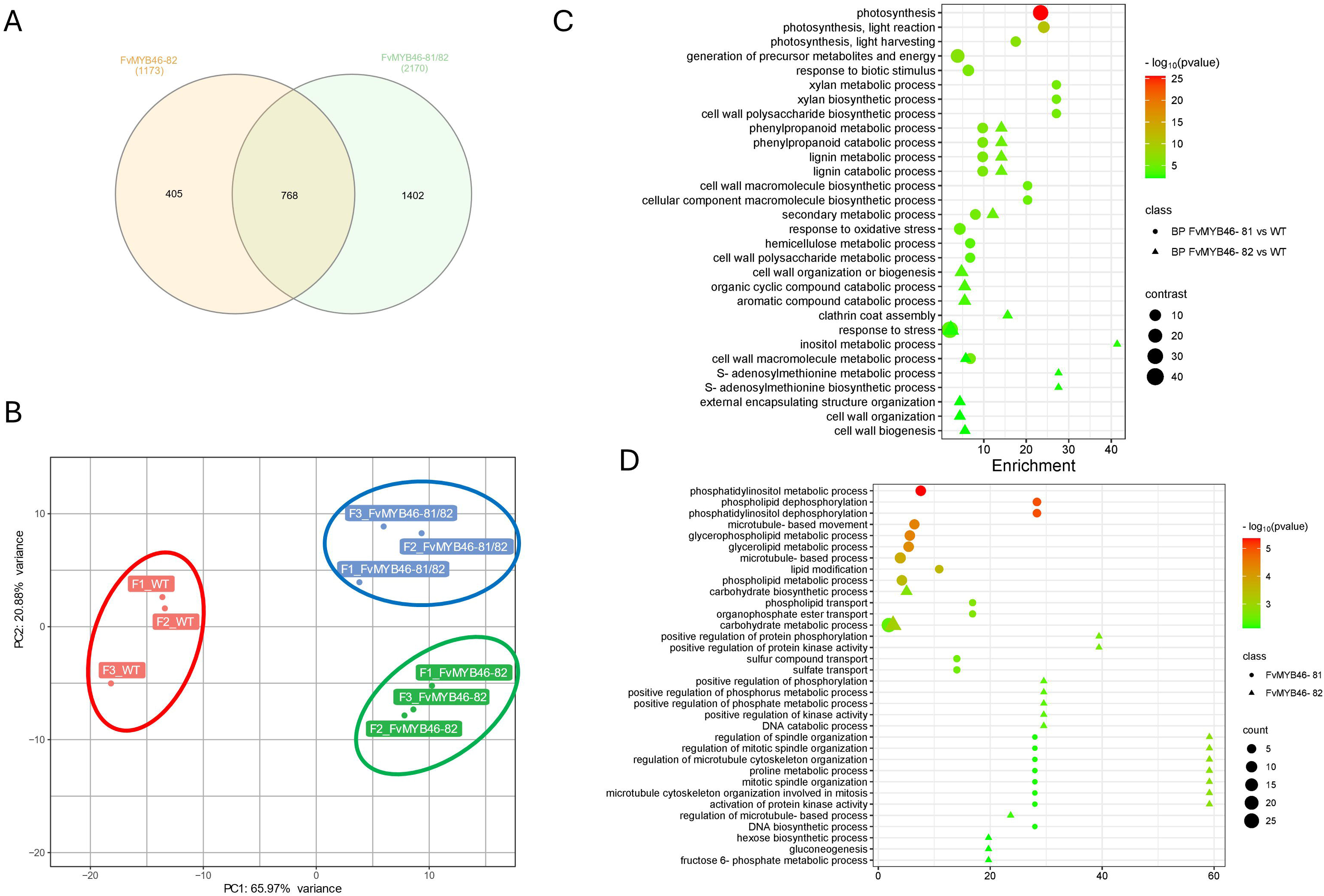
Co-expressed and unique DEGs in flowers FvMYB46-82 and FvMYB46-81/82 plants. (A) Venn diagram of co-expressed and uniquely expressed genes (FDR <0,05) after pairwise comparison of DEGs. B) Principal component analysis (PCA) analyses of samples based on RNA-Seq data. Pathway-enrichment bubble plots comparing enriched GO-terms (P < 0.01) in the biological process category for FvMYB86-81/82 vs wt (circles) and FvMYB46-82 vs wt (triangles) for downregulated DEGs (C) or upregulated DEGs (D). X-axis shows fold enrichment values.

### Enrichment analysis of functional terms in DEGs

GO (Gene Ontology) enrichment analysis of downregulated DEGs (P-value <0.01) in the biological process category for *FvMYB46-82* flowers, showed that terms related to secondary metabolites, cell wall, lignin biosynthesis and metabolism, and phenylpropanoid biosynthesis and metabolism were among the most enriched (Table 1). The most enriched GO terms in the biological process category of downregulated DEGs for *FvMYB46-81/82* flowers (Figure 3 c, Supplementary table 2) where similar to *FvMYB46-82*, except that the GO terms ‘photosynthesis’, ‘xylan biosynthesis’ and ‘xylan metabolic process’ also were enriched. For upregulated genes the terms ‘phosphatidylinositol metabolic process’ and ‘phosphatidylinositol dephosphorylation’ were enriched for *FvMYB46-81/82* flowers, while ‘carbohydrate metabolism’ and terms related to microtubule were enriched in both mutants (Figure 3 d, Supplementary table 3).

**Table 1.** Selected differently expressed genes in flowers and young berries of FvMYB46-deletion mutants compared to wild type, NS: Not statistically different, FDR <0,01.

|  |  |  | Flower |  |
| --- | --- | --- | --- | --- |
| Gene ID | Gene Name | Gene bank annotation | log2 FC | log2 FC |
|  |  |  | FvMYB46-82 | FvMYB46-81 |
| Signalling and transcription factors |  |  |  |  |
| FvH4_3g28890 | Myb46 | Myb Transcription Factor | -1,16 | -0,54 |
| FvH4_7g01020 | Myb83 | Myb Transcription Factor | NS | NS |
| FvH4_3g45450 | MYB5 | Transcription Repressor Myb5 | NS | -0,14 |
| FvH4_3g34960 | MYB59 | Transcription Factor Myb59 | NS | 0,22 |
| FvH4_5g17111 | MYB3-like | Transcription Factor Myb3-Like | -0,68 | -0,65 |
| FvH4_7g08972 |  | L10-Interacting Myb Domain-Containing Protein-Like | 9,28 | 9,23 |
| FvH4_6g33050 | NAC37 | NAC domain-containing protein 37 | 6,09 | 6,094 |
| Phenylpropanoid pathway |  |  |  |  |
| FvH4_6g16060 | PAL1 | Phenylalanine Ammonia-Lyase | NS | NS |
| FvH4_7g19130 | PAL2 | Phenylalanine Ammonia-Lyase | -0,55 | -0,86 |
| FvH4_2g05780 | CCOAMOT1 | Caffeoyl-Coa O-Methyltransferase | -0,81 | -1,07 |
| FvH4_3g06690 |  | Shikimate O-Hydroxycinnamoyltransferase-Like | NS | -0,75 |
| FvH4_6g28680 | CCR1 | Cinnamoyl-Coa Reductase 1-Like | -0,38 | -0,57 |
| FvH4_6g27940 | 4CL1 | 4-Coumarate--Coa Ligase 1-Like | -0,85 | -1,09 |
| Flavonoid, anthocyanin and lignin biosynthesis |  |  |  |  |
| FvH4_7g32990 | COMT1 | Caffeic Acid 3-O-Methyltransferase | NS | -0,43 |
| FvH4_7g25890 | CHI3 | Chalcone-Flavanone Isomerase 3 | -0,56 | -0,82 |
| FvH4_7g20870 | FvCHI1 | Chalcone-Flavanone Isomerase 1 | -0,58 | -0,79 |
| FvH4_7g33840 | FvUGT | Anthocyanidin 3-O-Glucosyltransferase 2 | -0,95 | -1,18 |
| FvH4_5g01170 | FvANS | Anthocyanidin Synthase | -0,84 | -1,04 |
| FvH4_7g01160 | FvCHS1 | Chalcone Synthase | -0,72 | -0,96 |
| FvH4_2g39520 | FvDFR | Dihydroflavonol 4-Reductase | -0,61 | -0,91 |
| FvH4_1g11810 | F3H | Flavanone-3-Hydroxylase | -0,49 | -0,69 |
| FvH4_3g40570 | C4H/CA4H | Trans-Cinnamate 4-Monooxygenase | -0,46 | -0,88 |
| fvh4_3g13000 |  | Anthocyanidin 3-O-Glucosyltransferase 7-Like | -2,01 | -2,22 |
| FvH4_3g02980 | FvANR | Anthocyanidin Reductase | NS | -0,73 |
| FvH4_2g39620 | EGS1 | Isoflavone Reductase-Like Protein/Eugenol Synthase | -0,53 | -0,77 |
| FvH4_1g29330 | C3H | p-coumaroyl-shikimate/quinic 3-hydroxylase | -0,472 | -0,525 |
| FvH4_6g30610 | CSE | caffeoyl shikimate esterase | -0,405 | -0,597 |
| Cell wall associated genes |  |  |  |  |
| FvH4_6g16911 | FvEXPA18 | Expansin-A9-Like | 2,45 | 2,68 |
| FvH4_3g36410 | FvEXPA9 | Expansin A6 | -0,67 | -0,68 |
| FvH4_6g15450 | IRX9 | Probable Beta-1,4-Xylosyltransferase Irx9 | NS | -0,62 |
| FvH4_1g00670 | CESA7/IRX3 | Cellulose Synthase A Catalytic Subunit 7 | -0,53 | -0,82 |
| FvH4_3g07420 | CESA8/IRX1 | Cellulose Synthase A Catalytic Subunit 8 | -0,28 | -0,58 |
| FvH4_3g38140 | Lac4/IRX12 | Laccase-4 | -0,81 | -1,06 |
| FvH4_6g11960 | Lac4/IRX12 | Laccase-4 | -0,53 | -0,98 |
| FvH4_4g19370 | Chl2 | Chitinase-Like Protein 2 | -0,83 | -1,29 |
| Carbohydrate metabolism |  |  |  |  |
| FvH4_4g15170 | FvSTP20 | Sugar transporter | 2,548 | 2,875 |
| FvH4_4g15160 | FvSTP18 | Sugar transporter | 2,388 | 2,434 |
| FvH4_4g15172 | FvSTP14 | Sugar transporter | 1,264 | 1,921 |
| FvH4_6g19270 | FvINV1 | Putative Beta-Fructofuranosidase/Cell wall invertase | 9,64 | 8,78 |
| FvH4_6g19271 | FvINV3 | Putative Beta-Fructofuranosidase/Cell wall invertase | 6,25 | 5,57 |
| FvH4_1g28890 | FvFRK3 | Fructokinase-6, chloroplastic | -0,40 | -0,53 |
| FvH4_2g00080 | FvFRK5 | Fructokinase-5 | -0,94 | -1,03 |

For the molecular function category, the most enriched GO terms for downregulated DEGs in flowers of both mutants were related to ‘oxidoreductase activity’. For category cellular component, ‘apoplast’ and ‘extracellular region’ were the most enriched categories in both *FvMYB46-81/82* and *FvMYB46-82* flowers.

A more detailed functional categorization of DEGs and associated metabolic pathways was done with Mercator enrichment and Mapman4 analysis (Figure 4 a-c, Figure S4, S5). This analysis confirmed the significant downregulation of genes involved in cell wall organization, lignin and monolignol conjugation and polymerization in flowers of both mutants compared to wild type. It also showed significant enrichment of genes involved in secondary metabolites metabolism, such as phenolics and flavone and flavonoid biosynthesis, and also showed significant enrichment of genes involved in redox homeostasis, especially glutathione S-transferase activities among the downregulated genes, while pectin and other cell wall proteins were upregulated.

**Figure 4.**
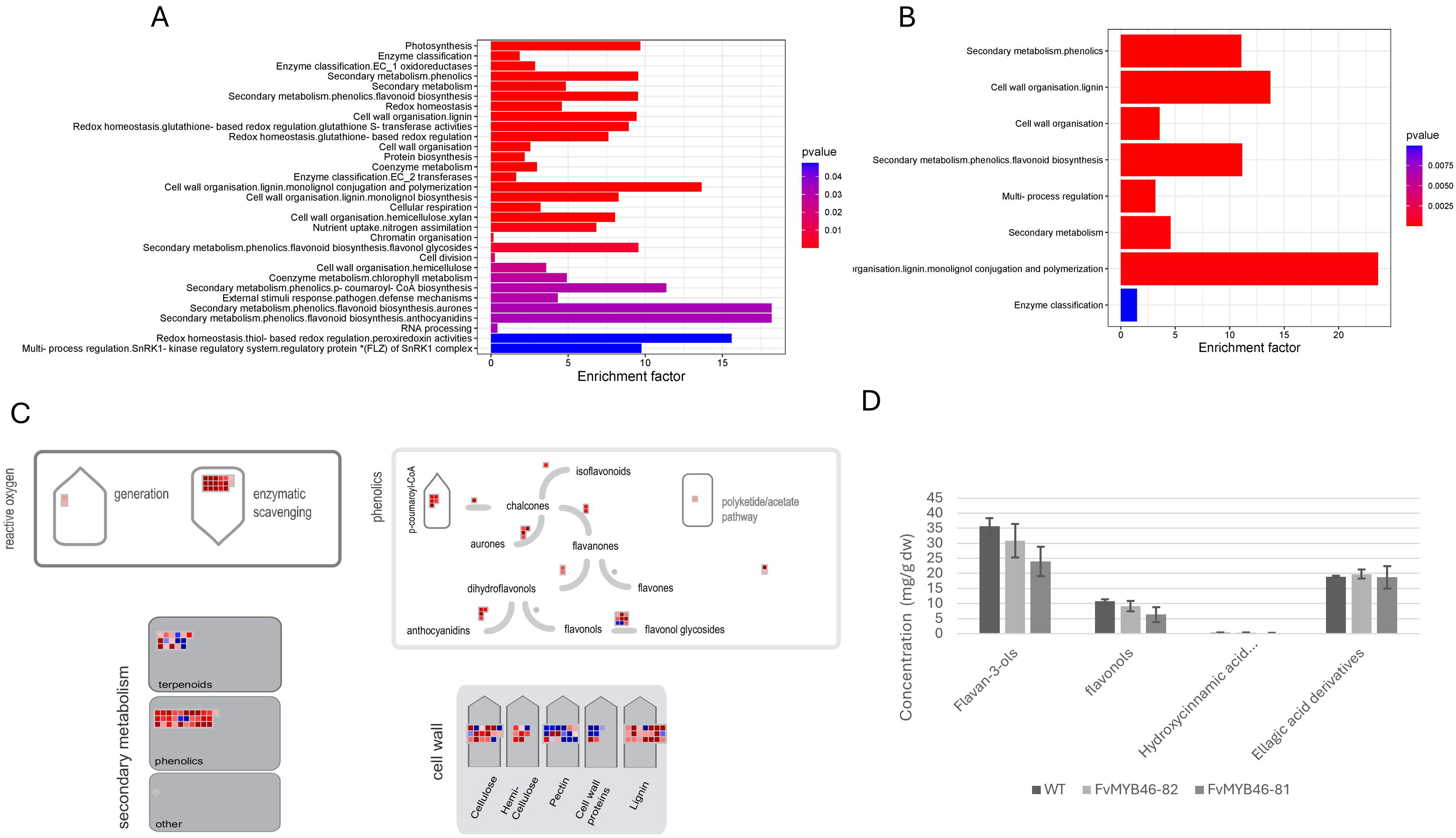
Mercator and Mapman visualization of enriched metabolic pathways in FvMYB46 mutants. A) Mercator enrichment analysis of differentially expressed genes (DEGs) for FvMYB46-81/82 (A) and FvMYB46-82 (B) mutant plants compared to wild type. C) Genes differently expressed in FvMYB46-81/82-mutants displayed onto metabolic pathways using the MAPMAN software: Secondary metabolism, including phenolics and terpenoids; flavonoid pathway; Redox homeostasis; and cell wall. Blue cells: upregulation in FvMYB46-82 compared to wild type; red cells: downregulation in FvMYB46-82 compared to wild type. D) Concentration of flavan 3-ols, flavonols, ellagic acid derivates and hydroxycinnamic acid derivatesin flowers using HPLC in mg/g dry weight.

### Analysis of phenolic compounds

HPLC-analysis of flower tissue identified 18 phenolic compounds that were grouped into flavonols, flavan-3-ols and derivatives of ellagic and hydroxycinnamic acids. The flavan-3-ols and flavonols, were reduced for both mutants compared to wild type, although the lowest concentration was detected in the *FvMYB46-81/82* mutant (Figure 4 d). Of individual compounds, epicatechin and proanthocyanin 3 belonging to the flavon-3-ols group, and quercetin-glucoside and Kaempferol-3-coumaroylhexoside, belonging to the flavonol group, were most downregulated (Figure S6). The concentrations of hydroxycinnamic acid derivatives were relatively low, with minor differences between the genotypes, while no significant differences were detected for ellagic acid derivatives.

## Discussion

### MYB46 regulates fruit setting rate

The R2R3 MYB transcription factor MYB46 is a central regulator of lignin and flavonoid biosynthesis and secondary cell wall formation in xylem vessels and fibers, but little is known about its activity in flowers and berries in *F. vesca*. In this study we used CRISPR-Cas9 constructs with an endogenous FvU6-1 promoter to introduce in-frame gene deletions (*FvMYB46-81*) or out of frame (*FvMYB46-82)* knock outs of FvMYB46. In contrast to overexpression-experiments where constitutive expression can cause unspecific ectopic effects, CRISPR-Cas9 editing makes it possible to study gene function in its native context.

The reduced pollen germination and fruit setting causing high frequency of berries arrested at the receptacle stage in *MYB46-81/82* and *MYB46-82* plants are similar to *F. vesca* plants with a CRISPR-cas9 mediated knock-out of the MADS box gene FveAGL62 which is required for early fruit development (L. Guo et al., 2022). This suggests that FvMYB46 regulates fertility and early fruit development in *F. vesca* corresponding to the specific expression in anthers and carpel walls. Before fertilization *FvMYB46* is expressed in flowers, specifically in anthers at flower development stage 11. At this stage the filaments reach its final size and the endothecium cells of the anther increase in size due to formation of lignified, cellulosic secondary wall thickenings necessary for anther dehiscence and pollen dispersal (Dawson et al., 1999; Hollender et al., 2012). In *Arabidopsis*, both repression and overexpression of MYB46 is associated with sterility due to anther indehiscence and ectopic secondary cell wall formation in stamens and carpels, respectively (Nguyen et al., 2019; Zhong et al., 2007).

Recently overexpression of FvNST1b, the closest homologue to AtNST1 which directly regulates MYB46 and anther dehiscence and fertility in *Arabidopsis*, was shown to promote secondary cell wall in various tissues including anthers and ovules in *F. vesca* (Dang et al., 2022). Like its RiMYB46 homologue, *FvMYB46* expression in the carpel walls of the green berries coincides with the activation of cell wall synthesis genes and extensive lignification when the carpels develop into dry achenes post fertilization (Zhou et al., 2023). Thus, the enrichment of the GO terms ‘phenylpropanoid metabolic and catabolic process’ and ‘lignin metabolic and catabolic process’ and “secondary cell wall biosynthesis” in downregulated DEGs in mutant-flowers suggest that the reduced pollen viability, ineffective fertilization and fruit set observed in *FvMYB46-81/82* and *FvMYB46-82* plants is caused by deregulated lignification and secondary cell wall formation in anthers or carpels. However, we did not observe any effect on total lignin content, although we observed a modest reduction in lignin content in endothecium cells facing the connective tissue of the filament suggesting that FvMYB46 positively regulates lignification of secondary cell walls in these tissues. Pollen development and dehiscence is tightly controlled, and both premature or delayed processes can cause sterility. The significant reduction in fruit set of *FvMYB46*-deletion mutants, although the effect on pollen germination and anther lignification were modest, suggest that FvMYB46 activity might be important for the correct timing of these processes to ensure efficient fruit set. The modest effect on lignin content of the *FvMYB46* deletions at the metabolite level were also reflected at the transcriptional level. Although there was a significant enrichment of lignin biosynthetic genes in DEGs downregulated in flowers, the fold reduction for the individual genes were modest. For example, several laccases homologous to Lac4 and Lac17 which are involved in lignin biosynthesis and degradation in *Arabidopsis* (Zhao et al., 2013), are weakly but significantly downregulated in *FvMYB46-81* and *FvMYB46-82* flowers (Table 1).

### MYB46 regulates flavonoid biosynthesis and stress signaling genes in fertilization and early fruit development

Several studies have shown that MYB46 enhances stress tolerance. In Apple and birch, up-regulation of MYB46 enhance drought and salt tolerance by regulating secondary cell wall biosynthesis and reduce reactive oxygen species by inducing genes encoding ROS scavenging enzymes (Chen et al., 2019; Geng et al., 2018; H. Guo et al., 2017). Terms related to secondary metabolite biosynthesis encompassing phenylpropanoid biosynthesis, especially flavonoid biosynthesis, as well as genes involved in enzymatic scavenging of reactive oxygen species were enriched in FvMYB46-mutants. In *F x ananassa* the general phenylpropanoid pathway and flavonoid biosynthesis enzymes CHS, PAL2, CHI, EGS1, CCoAOMT and F3H are expressed in flowers and early fruit development (Aragüez et al., 2013; Baldi et al., 2018). Consistent with the suggested role for FvMYB46 in regulating phenylpropanoid biosynthesis in early fruit development, these genes were downregulated in the *FvMYB46* mutants.

Silencing of chalcone synthase which is the first step in flavonoid biosynthesis, leads to impaired pollen tube growth in tomato (Schijlen et al., 2007) and reduced levels of anthocyanins, flavonols, and proanthocyanidins in *F x ananassa* (Lunkenbein et al., 2006). Metabolite profiling confirmed that the level of the flavonols, quercetin-glucoside and kaempferol-3-coumaroylhexoside, and the flavan-3-ols proanthocyanin 3 and epicatechin were reduced in open flowers of *FvMYB46* mutants. This suggests that FvMYB46 regulates the flavonoid biosynthesis and activity of ROS signaling genes in fertilization and early fruit development, most likely in the anthers and carpel walls, thus affecting the observed fruit set phenotype. Flavonoids are important for stress tolerance, pollen development, pollen viability, pollen germination. and pollen tube growth and thus have an important role in plant reproduction (Ylstra et al., 1994). Several studies in different plants show that abiotic stress and mutations in genes in the flavonoid biosynthesis pathway, including flavones and anthocyanins, impair pollen development and fertility. In tomato, reduced flavonol content in pollen and pollen tubes of the *anthocyanin reduced (are)* mutant lacking a functional flavonol 3-hydroxylae (F3H), were associated with reduced number of viable pollen grains, slower pollen tube growth and reduced pollen germination rate and higher abundance of total ROS and H2O2 under normal and heat stress conditions, consistent with flavonol controlling pollen development and pollen tube growth by scavenging reactive oxygen species (ROS) (Muhlemann et al., 2018). The *are* mutant also had smaller fruits and lower seed set compared to wild type. Interestingly, glutathione-S-transferases are enriched in the downregulated genes in the *FvMYB46* mutant and are co-expressed with *FvMYB46* in the carpel wall/ovary wall of the developing embryo. Glutathione-S-transferases are detoxifying and ROS scavenging enzymes improving the plants resistance to stress. Thus, increased stress susceptibility or impaired ROS signaling may be influencing the reduced fruit set phenotype observed in *FvMYB46*-mutants.

Several other genes important for successful fertilization were differently expressed in the *FvMYB46*-mutants. For example, expansins which are proteins involved in loosening cell walls, were differently regulated in the FvMYB46 flowers. Expansins are also believed to be involved in softening cell walls of the stigma, facilitating growth and penetration of the pollen tube and successful fertilization (Marowa et al., 2016; Mollet et al., 2013). In both FvMYB46-deletion mutants the GO term ‘carbohydrate metabolism’ was enriched for upregulated genes. The cell wall invertase homologues *FvINV1* and *FvINV3* were strongly induced in the *FvMYB46* flowers. Cell wall invertases (CWI) hydrolyses sugars to fructose and glucose in the extracellular matrix for further sugar processing and starch accumulation in anthers and ovaries (De Storme & Geelen, 2014). CWI’s play important roles in reproduction, and downregulation or knock-out is associated with reduced pollen germination, ovary activity and fruit set (Ezura et al., 2023). In contrast, increased CWI activity is associated with sustained fruit set, cell division and fruit growth possibly due to reduced PCD. CWIs are generally coregulated with sugar transporter proteins which is required for efficient transport of glucose and fructose from phloem to parenchyma cells. Fructokinases are then required to phosphorylate fructose to fructose-6-phosphate which is crucial for efficient fruit set by facilitating unidirectional flux from sugar metabolism to the generation of cell wall components and energy (Ezura et al., 2023; Shinozaki et al., 2020). In *FvMYB46* mutant flowers, we observed 3 sugar transporters that were coregulated with the invertases, while several fructokinases and cellulose synthase (CesA) genes and genes involved in xylan biosynthesis were downregulated (Table 1). Thus, although high activity of invertases and sugar transporters are associated with increased fruit set, the concomitant downregulation of fructokinases and cell wall genes might cause reversed carbon flux having negative impact on fruit set in the *FvMYB46*-mutants. Thus, we suggest that FvMYB46 is involved in sugar signaling in reproductive organs required for efficient fruit set in *Fragaria vesca*.

### Redundancy and dominant negative effect

The low or missing effect on level lignin content in *FvMYB46* deletion lines is consistent with the low increase in total lignin content using MYB46 overexpression constructs in both *Arabidopsis* and *F*. *vesca* (Kim et al., 2014; Zhang et al., 2023). Weak effect on lignin level in *FvMYB46* deletion mutants might be due to redundancy with the endogenous paralogue FvMYB83 transcription factor, as is seen in *Arabidopsis* (McCarthy et al., 2009; Zhong et al., 2007), and which were co-expressed with *FvMYB46* in flowers and early fruit development. However, the observed fruit setting phenotype and significant effect on different biosynthetic pathways, suggest that FvMYB83 or other potential homologs are not completely redundant with FvMYB46 in flowers possibly partly because FvMYB83 has a lower expression than MYB46. Interestingly, we observed a stronger effect on pollen germination and a larger number of DEGs with a stronger up- or downregulation in bi-allelic *FvMYB46-81/82* flowers compared to *FvMYB46-82*. This suggests that the expressed in-frame gene product *FvMYB46-81* with a deletion in the conserved first MYB DNA binding domain, can act as a dominant negative, for example by competing with MYB83 by binding a non-functional protein to the native target promoters or by interaction with other proteins. A similar dominant negative effect was seen for MYB46 in *Arabidopsis* (Zhong et al., 2007), however in contrast to this study where the MYB46 protein were ectopically expressed by the constitutive 35S-promoter, the expression of the putative deletion protein is driven by the endogenous promoter. Thus the phenotype is more likely reflecting the effect of the native protein competing with redundant proteins in its natural context and location. At the transcriptomic level, there were a significant number of overlapping DEGs between the mutants, although there were almost twice as many in the *FvMYB-81/82* mutant. The majority of GO-terms were similar for both mutants, although some were unique for the *FvMYB-81/82*-bp mutant like ‘photosynthesis’ in downregulated DEGs and terms related to phosphatidylinositol in upregulated DEGs.

However, although these DEGs and the stronger up or downregulation of similar DEGs in *FvMYB46-81/82* flowers, likely can explain the slightly different effect on production of phenolic compounds and pollen germination between the mutants, they are not essential for the major fruit set phenotype observed in both mutants. To summarize, we have shown that CRISPR-Cas9 mediated deletions of FvMYB46 caused reduced pollen germination and fruit set compared to wild type *F. vesca*. Transcriptional profiling showed that genes involved in secondary cell wall formation, lignin and flavonoid biosynthesis, pollen tube growth, and scavenging of reactive oxygen species were enriched in downregulated genes, while genes involved in carbohydrate metabolism were upregulated in *FvMYB46*-flowers. The role in flavonoid biosynthesis was supported by metabolite profiling which demonstrated a reduction of flavonols and flavan-3-ols in mutant flowers. FvMYB46 expression coincides with secondary cell wall formation and important developmental processes in anthers and carpels important for pollen viability and anther dehiscence before fertilization, and early fruit development after fertilization. Thus, our work demonstrates that FvMYB46 control efficient fertilization and fruit set by biosynthesis of flavonoids and cell wall components and regulating genes involved in ROS signaling and carbohydrate metabolism in *F. vesca* reproductive organs.

## 4. Methods

### Construction of CRISPR MYB46 knock-out plasmids

For stable transformation experiments for knock-out of MYB46, we synthesized a construct containing two individual expression cassettes containing FvU6-1-gRNA1-scaffold-terminator, and FvU6-1-gRNA2-scaffold-terminator (Supplementary material), respectively with GenArt (Thermo Fisher). The fragment was then cloned into the pCAS9-TPC vector (Fauser et al., 2014) using PacI giving pCAS9-TPC/MYB46_2XgRNA construct used for *Agrobacterium* mediated transformation of *F. vesca* Hawaii-4. The sgRNA1 (TGAGTAGTAATAATAAGCTG) and sgRNA2 (CCAAGGCTGCTGGAGCGACG) sequences targeting the first exon of *FvMYB46* were designed using the CRISPR-P 2.0 program (H. Liu et al., 2017). CRISPR-P 2.0 was also used to predict potential off-targets for gRNA1 and 2 which were amplified using specific primers (Supplementary Table Y) and sequenced.

### Agrobacterium mediated stable transformation

Leaf discs of *F. vesca* H4X4 were transformed using the pCAS9-TPC/MYB46_2XgRNA construct in *Agrobacterium tumefaciens* strain GV3101. *F. vesca* Hawaii 4 seeds were germinated on ½ MS (Murashige & Skoog) medium for 30 days. From *in vitro* plants, petioles and single leaves were cut into 1-2 cm pieces dipped in resuspended *Agrobacterium* pellet in co-cultivation medium containing MS medium, 2% sucrose, pH 5.8, and freshly added 100µM acetosyringone. Cut leaf discs were incubated in co-cultivation media for 60 minutes at room temperature. Afterwards leaf discs were dried and moved to MS callusing media containing 2% sucrose, pH 5.8, 3 mg/L BAP, 0.2 mg/L IBA. Putative transformants were selected on MS callusing media containing 2% sucrose, pH 5.8, 3 mg/L BAP, 0.2 mg/L IBA and, 3 mg/L BASTA. After 10 weeks, calluses were transferred to shoot inducing media containing MS media with 2% sucrose, pH 5.8, 1mg/ml BAP and, 0.2 mg/ml IBA.

Developed shoots were transferred to MS rooting media with 1% sucrose, pH 5.8 and, BASTA selection. Plants of around 5 cm height were moved to soil and maintained in the growth room.

### Plant growth and phenotyping

*F. vesca* ‘Hawaii-4’ plants were grown in topsoil in 400 mL pots cultivated in a growth room with 14 h light (∼100 mmol m-2 s-1 photosynthetically active radiation (PAR) at 24 °C) and 10 h darkness (19°C) at 40-45% relative humidity for phenotypic inspection and for production of material for all transcriptomic analyses For transformation with *Agrobacterium* seed produced in green house were sterilized and grown in vitro ½ MS medium in Petri dishes in plant growth room. For runner and seed production, plants were propagated in the greenhouse under standard conditions. Fruit set was determined by calculating the percentage of fruits fully developing to the ripe stage divided by the total number of open flowers. To ensure that all flowers were at the same developmental stage, they were marked at the pre-anthesis stage. Flowers from 12 plants of wild type, *FvMYB46-81* and *FvMYB46-82* plants were counted over 40 days, starting from day one. All the flowers bloomed in this period, flowers that resulted in aborted berries and flowers resulted in berries were quantified for 40 days. For pollen tube germination, we used culture medium for *in vitro* germination of strawberry pollen developed by Yamaguchi et al. (2024), without agar. Briefly, the flowers were collected from greenhouse in the morning, and the pollen were cultured into 10% sucrose and 0.1% boric acid medium. A total of 300µl of medium was dispensed onto a microscope slides glass cavity (with 14 mm diameter and 6.5 mm−^1^ curve surface), and the pollen were brushed into the medium. From each genotype, seven flowers (technical replicates) were collected, and 4 biological replicates were investigated in the study. The germination of the pollen was performed after incubation for 3 h at 22°C by counting 100 pollen grains from each slide using an optical microscope. The germination of the pollen was evaluated by measuring the diameter length of pollen, pollen that has longer diameter than the pollen grain, was accounted as germinated. The rate of pollen germination was calculated as published by Yamaguchi et al. (2024), the variance between the mutant genotypes and the wild type was tested using one way ANOVA, and significance test was performed using Tukey’s multiple-range test (P < 0.01 or 0.05) in R software (R Core Team 2024).

### DNA and RNA isolation

About 100 mg of different tissues were ground in liquid N2 and used for DNA extraction using the DNeasy Plant Mini Kit (Qiagen), according to the manufacturer’s instructions. Total RNA was isolated from 100 mg tissue using the Spectrum Plant Total RNA Kit with minor modifications (Badmi et al., 2022). Briefly, preheated lysis buffer containing CTAB (2%), PVPP (2%), Tris-Cl (pH 8.0, 100 mM), EDTA (pH 8.0, 25 mM), NaCl (1 M) and b-mercaptoethanol (1%) was added and mixed with 100 mg of tissue powder and incubated at 65 °C for 8 min with vortexing for the first 60 seconds. After centrifugation for 10 min at 13,000 rpm, the supernatant was mixed with an equal volume of chloroform:isoamyl alcohol (24:1) and centrifuged again for 10 min at 4 °C. The supernatant was transferred to the kit’s filtration column (blue retainer ring), and from this step we followed the manufacturer’s instructions. On-column DNase I treatment was done to ensure DNA-free total RNA.

### Quantitative real-time PCR

A total of 500 ng RNA from different tissues was used to synthesize cDNA using iScript cDNA Synthesis Kit (Biorad). Quantitative real-time PCR (qRT-PCR) was performed using the CFX96TM Real-Time System (Biorad), and the SsoAdvanced Universal SYBR Green Supermix (Biorad) with the gene specific primers in Supplementary table Y. The relative expression levels were determined using the comparative threshold cycle (ΔCq) method (W. Pfaffl, 2001). *EF1A* was used as an endogenous control gene to normalize the data (W. Pfaffl, 2001).

### Transcriptomic profiling using RNA sequencing

Transcriptomic profiling of *F. vesca* Hawaii-4 open flowers and unripe berries 7 DAP were performed by BGI Genomics Co., Ltd. Five flowers or berries from 5 different wild type and mutant plants were harvested together for each replicate. Three biological replicates each of wild type, FvMYB46-81 and FvMYB46-82 flowers were used for RNA isolation. RNA samples were sent to Beijing Genomics Institute (BGI, https://www.bgi.com), Hong Kong for cDNA library construction paired-end sequencing (PE100, 40M) and sequencing using a DNA nanoball sequencing (DNBSEQ™) technology. To produce clean and highly reliable data software SOAPnuke (Version: SOAPnuke.2.2.1 Parameters: −l 15 -q 0.5 -n 0.05 -i (https://github.com/BGI-flexlab/SOAPnuke)) was used to exclude low-quality readings, readings polluted by adapters, and readings with an excessive number of unknown bases.

Sequences were trimmed and quality checked using Trimmomatic (version 0.39 (Bolger, A. M., Lohse, M., & Usadel, B. (2014). Trimmomatic: A flexible trimmer for Illumina Sequence Data. Bioinformatics, btu170) using settings recommended for paired-end reads. An average of 21 million read-pairs for each sample were left after filtering, and these were mapped against v4.0.a2 of the *F.* vesca genome (Li et al., 2019) using the STAR aligner (version 2.7.11b) (Dobin et al., 2013). Additional functional annotation of genes was also downloaded for this version of the genome (https://www.rosaceae.org/species/fragaria_vesca/genome_v4.0.a2) The data were submitted to NCBI’s Sequence Read Archive (SRA) as submission SUB14564935.

In order to identify differentially expressed genes (DEGs), the R-wrapper SARTools (version 1.8.1) (ref - http://dx.doi.org/10.1371/journal.pone.0157022) was used with recommended settings (https://github.com/PF2-pasteur-fr/SARTools/blob/master/template_script_edgeR.r) to run edgeR (version 3.42.4) (ref - doi:10.1093/bioinformatics/btp616). DEGs were scored as significant when the false discovery rate (alpha) was less than 0.05.

### Functional categorization of DEGs using GO and Mapman enrichment analysis

Enriched GO-terms were identified using the R-package GOstats (Falcon S, Gentleman R (2007). “Using GOstats to test gene lists for GO term association.” Bioinformatics, 23(2), 257-8.) (version 2.66.0), and FDR cutoff of 0.01 using functional annotation of the *F. vesca* v4.0.a2 transcriptome (www.rosaceae.org). GO pathway enrichment plot was plotted with SRplot (www.bioinformatics.com.cn/en). The Mercator4 v6.0 online tool (https://www.plabipd.de/mercator_main.html) was used to functionally annotate and classify all the *F. vesca* transcripts into hierarchically structured bins and combined with DEG analysis and displayed onto metabolic pathways with the MAPMAN software version 3.7.0 (Schwacke et al., 2019; Thimm et al., 2004). DEGs with FDR <=0.01 were plotted in Venn diagrams using InteractiVenn (Heberle et al., 2015).

### Extraction of phenolic compounds and HPLC

Phenolic compounds were extracted from the samples as described by (Nybakken et al., 2018). A 5 mg aliquot of each sample was subjected to extraction using methanol (HPLC - gradient grade, VWR International LLC, Randor, USA). The extraction process involved homogenization with 10 ceramic balls at a velocity of 6.5 m/s for 30 seconds, utilizing a VWR Bead Mill MAX homogenizer (VWR International, LLC Radnor, PA, USA). Post-homogenization, samples were iced for a 15-minute duration before undergoing centrifugation at a speed of 15,000 rpm for 3 minutes, facilitated by an Eppendorf centrifuge 5417C. The supernatant was decanted into a 5 ml tube, and the remaining residue was re-dissolved in 400 μl of methanol followed by homogenization and centrifugation identical to the previous steps. This extraction process was reiterated twice more, pooling the supernatants each time. Subsequently, the pooled supernatants were subjected to evaporation in a vacuum centrifuge (Eppendorf concentrator plus) before the dried extracts were reconstituted in 500 μl of a 1:1 (v/v) mixture of methanol and water, assisted by an ultrasonic cleaner (mod. no. USC200TH; VWR International LLC, Randor, USA).

HPLC analysis was performed on the methodology by (Julkunen-Tiitto & Sorsa, 2001). The apparatus utilized was an Agilent 1200 series HPLC system, equipped with a binary pump and a diode array detector (DAD) system (Agilent Series 1200, Agilent Technologies, Waldbronn, Germany). Separation was conducted on a Thermo Scientific 50 × 4.6 mm ODS Hypersil column with a particle size of 3 μm (Thermo Fisher Scientific Inc., Waltham, USA), maintained at a temperature of 25 °C. The mobile phases consisted of 1.5% tetrahydrofuran and 0.25% acetic acid in ultrapure water (Mobile Phase A) and 100% MeOH (Mobile Phase B). The gradient for Mobile Phase A was as follows: 0–1.5 min, 100% A; 1.5–3 min, 100– 85% A; 3–6 min, 85–70% A; 6–12 min, 70–50% A; 12–20 min, 50% A; 20–22 min, 50–0% A. The flow rate was maintained at 2 ml min−1, and the injection volume was 20 μl. Compounds identification was achieved by analysis of retention times and UV spectra. The absorptions at 270, 320 and 360 nm, were used to calculate concentrations by comparing with commercial standards.

### Gene IDs and accession numbers

The nucleotide sequence of *Fragaria vesca* transcription factor *FvMYB46* has gene id FvH4_3g28890 and Genbank accession number XM_004291742.2.

## Acknowledgments

We thank Monica Skogen and Eva Grodås for technical help with experiments and Vinh H. Le for propagating plant material used in this work.

## Data Availability Statement

The data were submitted to NCBI’s Sequence Read Archive (SRA) as submission SUB14564935.

