## Supplementary figures for "CRISPR-Cas9 mediated deletions of *FvMYB46* reduces fruit set and biosynthesis of flavonoids in *Fragaria vesca*"

### Supplementary figure S1.

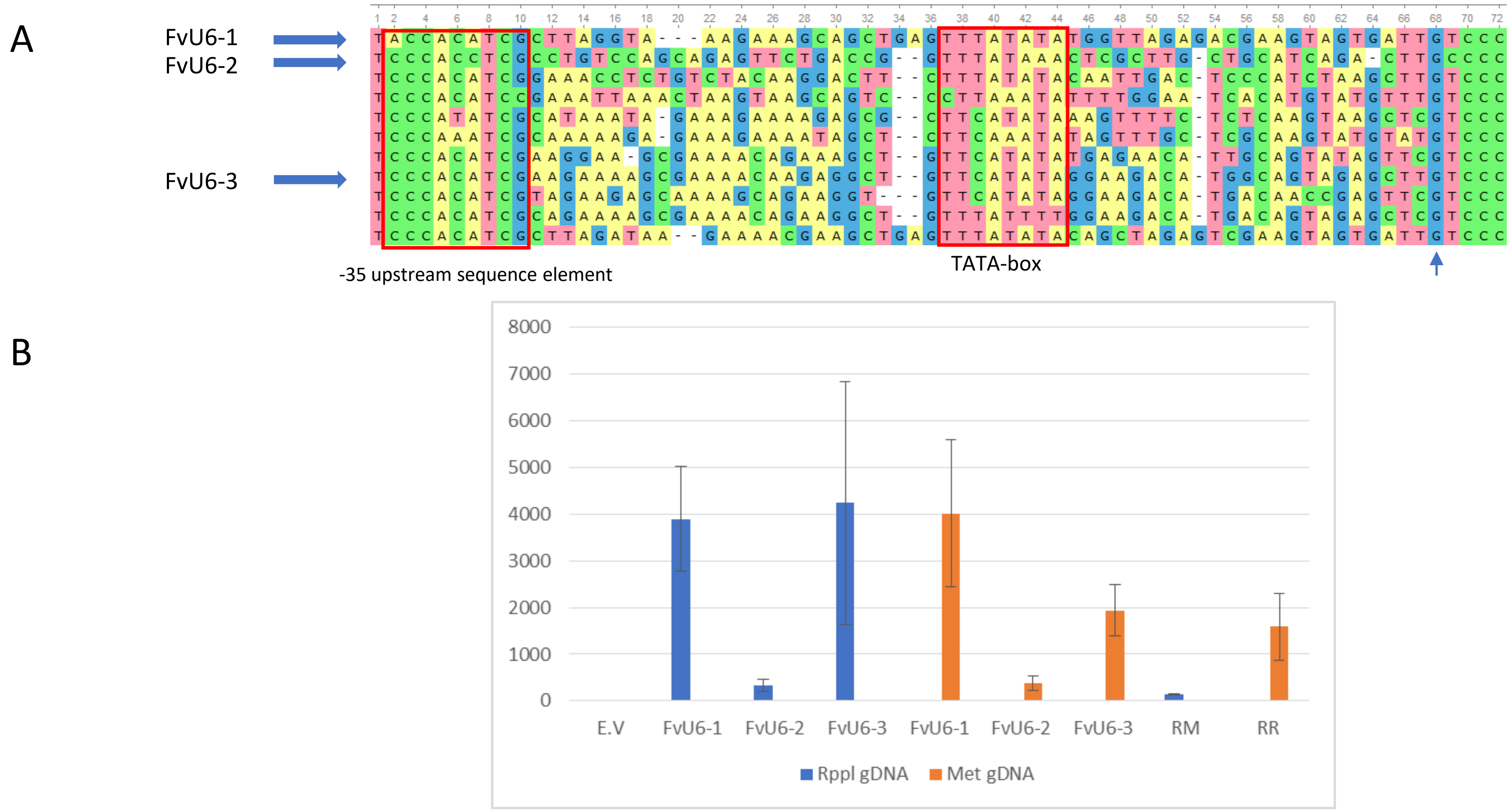


Supplementary figure 1. Identifying endogenous U6-promoters for gRNA expression *in F. vesca*. A) Alignment of the upstream U6-promoter sequence of snRNA genes identified in *F. vesca*. The promoters used for transient expression analyses in transient expression assays is indicated. Blue upwards arrow: transcriptional start site. B) Transient expression analysis of 2 different gRNA (RPPL1, Met1) in berries of *F. vesca* using the FvU6-1, Fv-U6-2 or Fv-U6-3 promoters compared to AtU6-promoter (RM, AtU6-Met1-gRNA; RR, AtU6-RPPL1), E.V, Empty vector;

### Supplementary figure S2

A

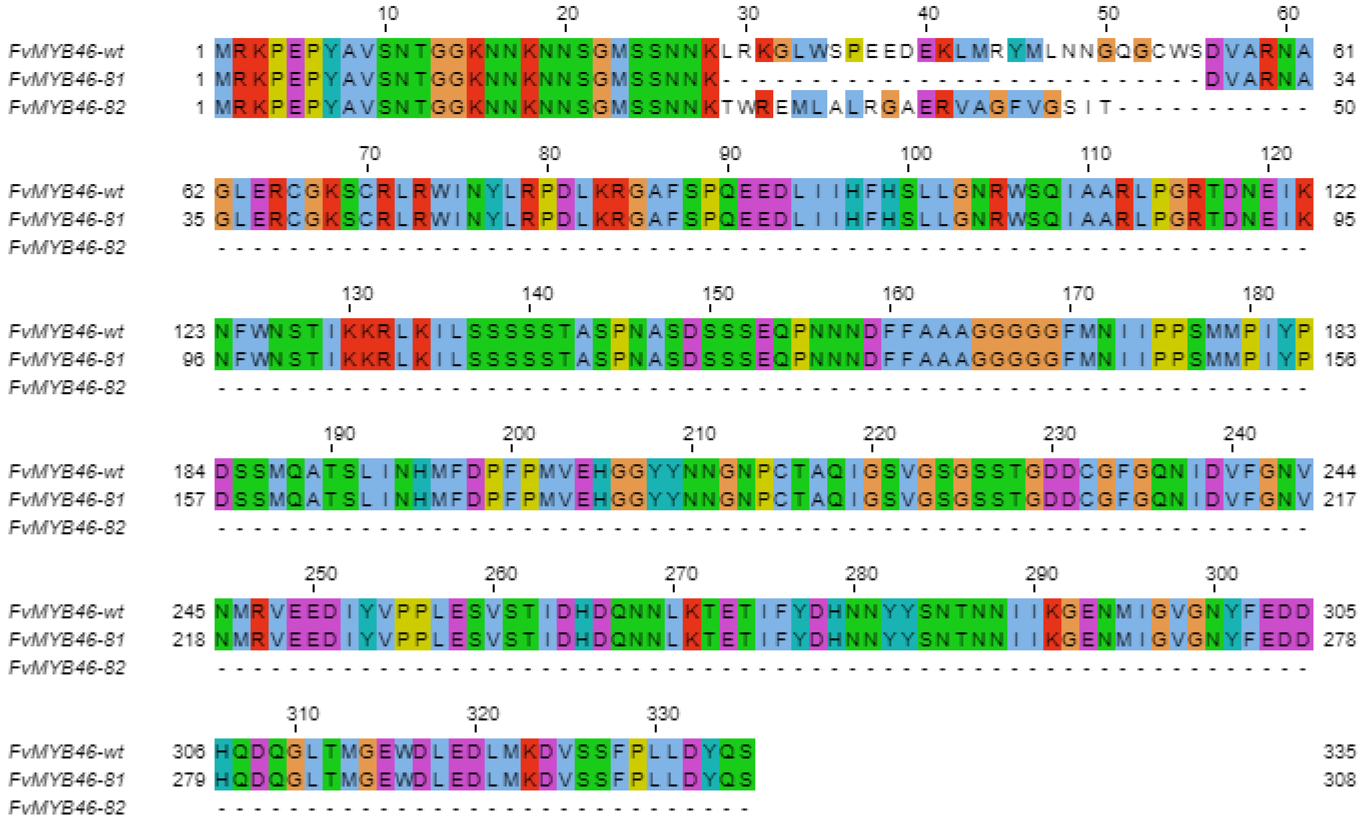


B

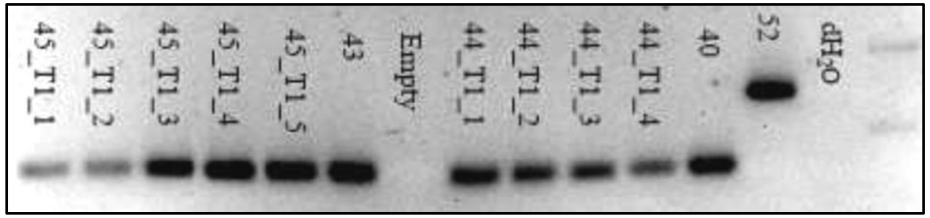


C

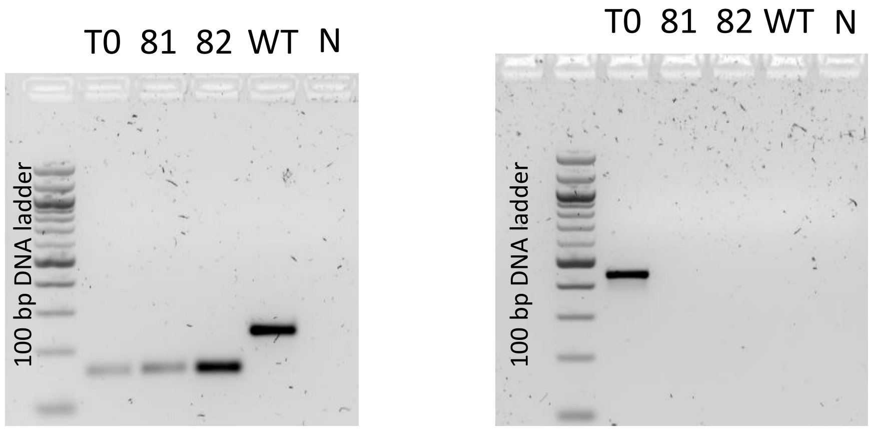


**Supplementary figure 2. CRISPR-cas9 mediated deletions of FvMYB46.** A) Alignment of predicted FvMYB46-81 and FvMYB46-82 translation products with wild type FvMYB46-protein. B) PCR-screening of DNA from T0-plants 40, 53 and 52 (neg) and progeny of 44 and 45-plants with gene specific primers flanking gRNA1 and gRNA2. Progeny of 43 and 44 were renamed FvMYB46-82 and FvMYB46-81/82 respectively C) Genotyping of DNA from T0 (44), FvMYB46-81/82, FvMYB46-82 and WT-plants used for transcriptomic and phenotypic studies (left). Genotyping of T0, FvMYB46-81/82 and FvMYB46-82 plants with Cas9-specific primers demonstrating no presense of T-DNA in homozygous and biallelic plants used in this study.

### Supplementary figure S3

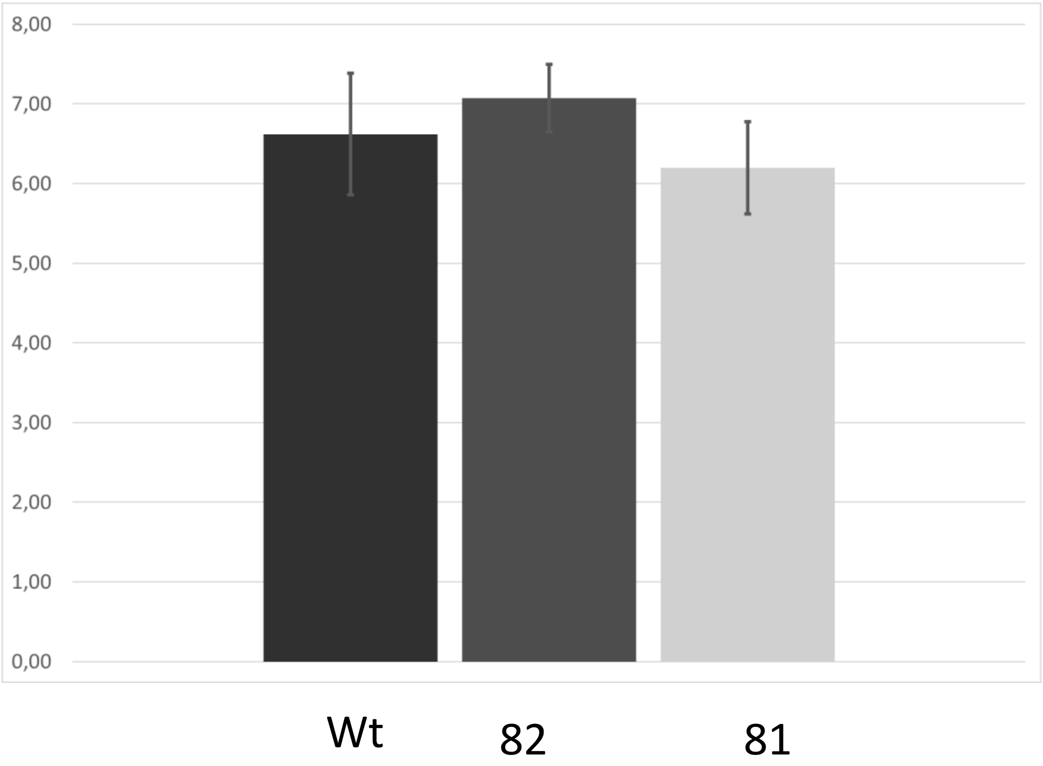


Supplementary figure 3. Lignin content in flowers of wt, FvMYB46-82 and FvMYB46-81 flowers determined by the Klason procedure

### Supplementary figure S4

A

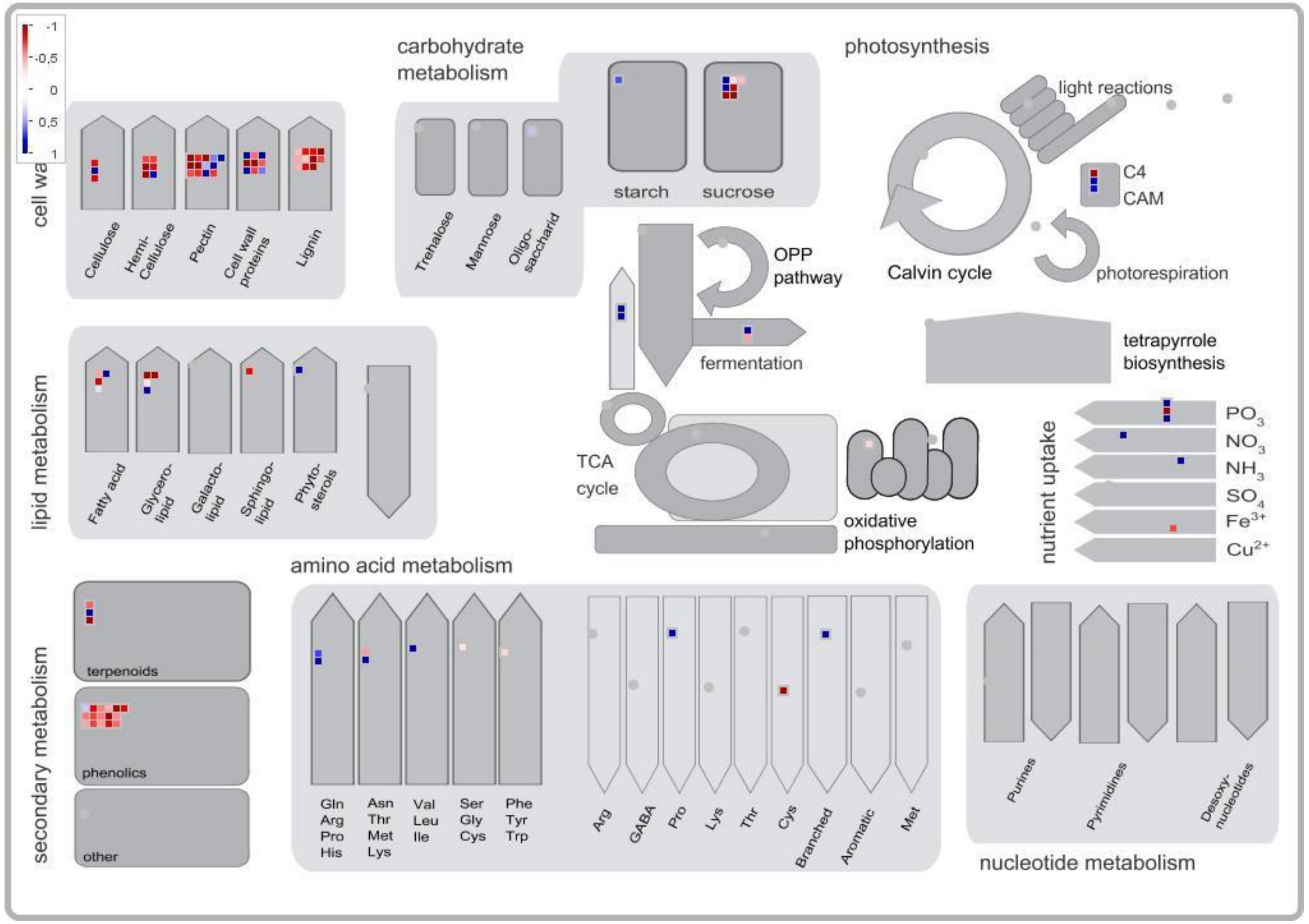


**Figure 5. Visualization of enriched metabolic pathways in the FvMYB46-82 mutant.** Differentially expressed genes (DEGs) in mutant plants compared to wild type displayed onto metabolic pathways using the MAPMAN software for flowers. Blue cells: upregulation in FvMYB46-82 compared to wild type; red cells: downregulation in FvMYB46-82 compared to wild type.

### Supplementary figure S5

A

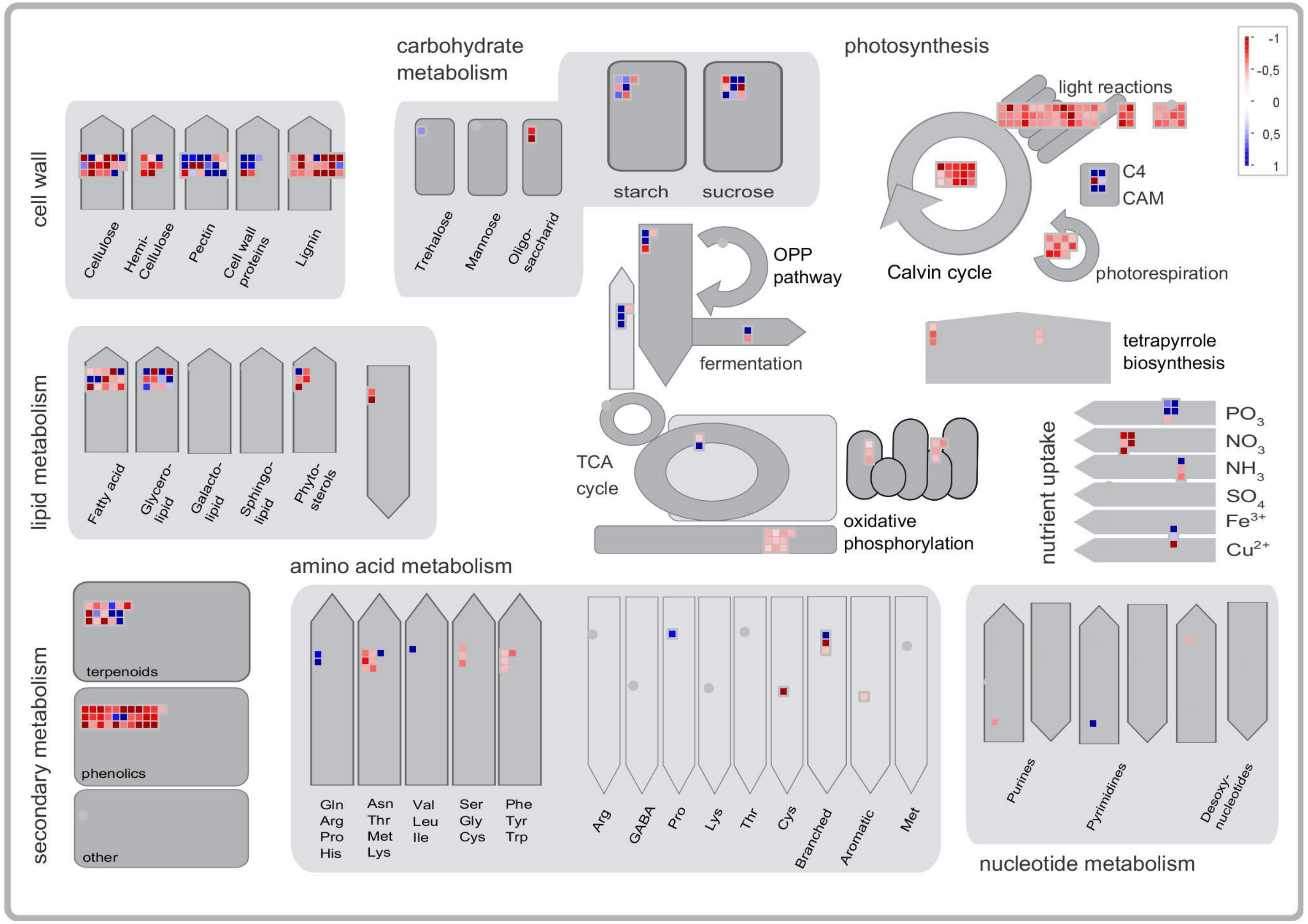


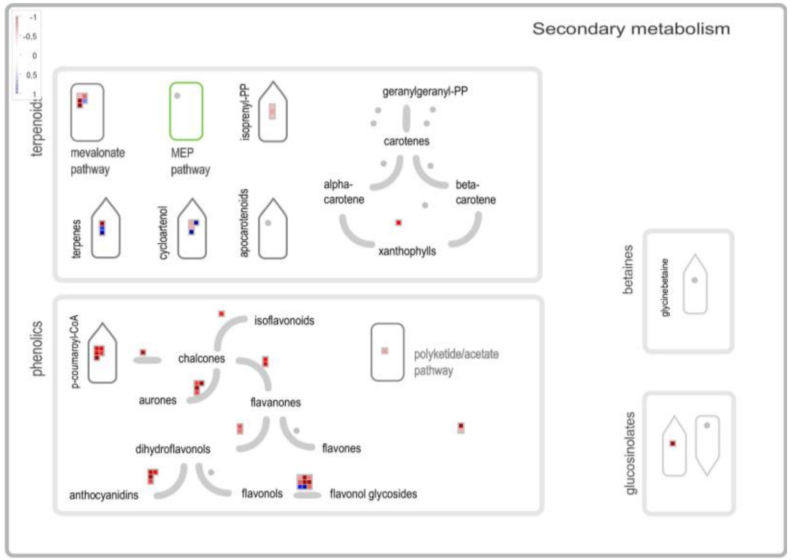


**Figure 5. Visualization of enriched metabolic pathways in the FvMYB46-81/82 mutant.** Differentially expressed genes (DEGs) in mutant flowers compared to wild type displayed onto metabolic pathways using the MAPMAN software. A subset of the data are shown in figure 5 C. A) Central metabolic pathways. B)) Secondary metabolism, including phenolics, flavonoid biosynthesis and terpenoids. Blue cells: upregulation in FvMYB46-81/82 compared to wild type; red cells: downregulation in FvMYB46-81/82 compared to wild type.

### Supplementary figure S6

#### Phenolic compounds analyzed in flowers

| COMPOUND | WILD-FLOWERS | 44-3-FLOWERS | 44-4-FLOWERS |
|---|---|---|---|
| CATECHIN | 0,86±0,09a | 0,94±0,33ab | 0,57±0,27abc |
| PROANTHOCYANIN 1 | 6,23±1,59a | 5,32±1,97b | 5,91±0,23b |
| EPICATECHIN | 5,17±0,29a | 2,19±0,27a | 2,69±0,88ab |
| PROANTHOCYANIN 2 | 3,1±0,7a | 2,69±0,44a | 2,09±0,76a |
| PROANTHOCYANIN 3 | 20,27±1,38a | 19,69±3,09a | 12,71±3,65b |
| SUM FLAVAN-3-OLS | 35,64±2,7a | 30,82±5,55ab | 23,98±4,92b |
| COUMARIC ACID-DERIVATIVE | 0,09±0,02a | 0,1±0a | 0,07±0,01ab |
| COUMARIC ACID-DERIVATIVE | 0,18±0,04a | 0,16±0,03a | 0,12±0,04a |
| COUMARIC ACID-DERIVATIVE | 0,07±0,02a | 0,07±0,01a | 0,05±0,01a |
| SUM HYDROXYCINNAMIC ACID DERIVATIVES | 0,34±0,04a | 0,32±0,04a | 0,24±0,07a |
| ELLAGIC ACID - DERIVATIVE 1 | 0,3±0,02a | 0,32±0,02a | 0,32±0,07a |
| ELLAGIC ACID - DERIVATIVE 2 | 0,42±0,06a | 0,35±0,04a | 0,21±0,08b |
| ELLAGIC ACID - DERIVATIVE 3 | 15,29±0,18a | 17±1,38a | 15,94±3,88a |
| ELLAGIC ACID - HEXOSIDE | 2,85±0,17a | 2,09±0,09ab | 2,25±0,48abc |
| SUM ELLAGIC ACID DERIVATIVES | 18,86±0,37a | 19,76±1,53a | 18,72±3,72a |
| QUERCETIN-3-GLUCURONIDE | 0,04±0a | 0,06±0,01ab | 0,04±0,01abc |
| QUERCETIN-GLUCOSIDE | 6,74±0,44a | 5,82±1,27ab | 3,87±1,4b |
| QUERCETIN-3-MALONYLGLUCOSIDE | 0,07±0,01a | 0,11±0,01ab | 0,09±0,02bc |
| KAEMFEROL-GLUCOSIDE | 0,24±0,01a | 0,19±0,04ab | 0,12±0,04bc |
| KAEMPFEROL-3-GLUCURONIDE | 2,51±0,13a | 2,13±0,36ab | 1,63±0,75b |
| KAEMPFEROL-3-HEXOSIDE | 0,32±0,02a | 0,29±0,03a | 0,22±0,08ab |
| KAEMPFEROL-3-COUMAROYLHEXOSIDE | 0,8±0,12a | 0,49±0,05b | 0,37±0,17bc |
| SUM FLAVONOLS | 10,72±0,66a | 9,09±1,7ab | 6,34±2,43b |

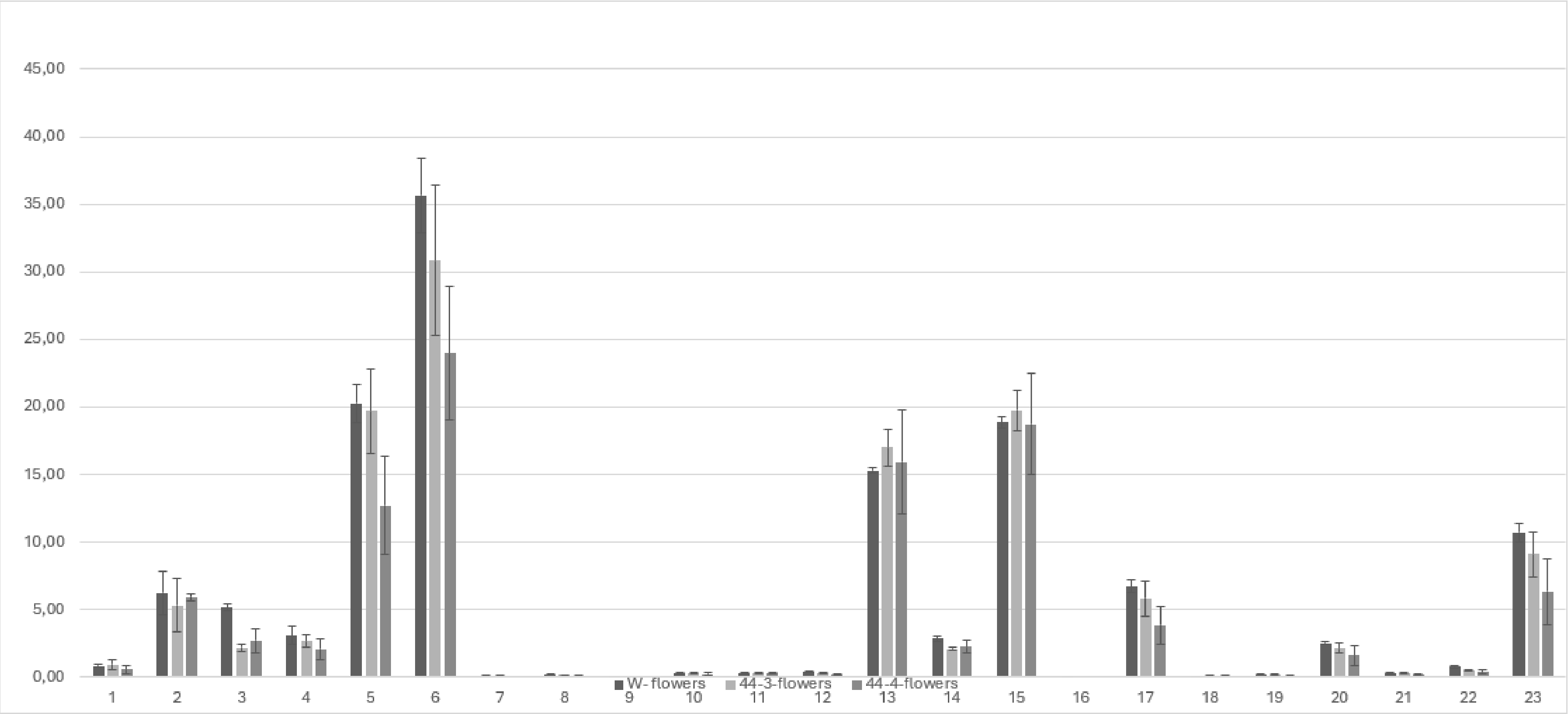


1 Catechin
2 Proanthocyanin 1
3 Epicatechin
4 Proanthocyanin 2
5 Proanthocyanin 3
6 Sum Flavan-3-ols
7 Coumaric acid-derivative
8 Coumaric acid-derivative
9 Coumaric acid-derivative
10 Sum Hydroxycinnamic acid derivatives
11 Ellagic acid - derivative 1
12 Ellagic acid - derivative 2
13 Ellagic acid - derivative 3
14 Ellagic acid - hexoside
15 Sum Ellagic acid derivatives
16 Quercetin-3-glucuronide
17 Quercetin-glucoside
18 Quercetin-3-malonylglucoside
19 Kaemferol-glucoside
20 kaempferol-3-glucuronide
21 Kaempferol-3-hexoside
22 Kaempferol-3-coumaroylhexoside
23 Sum flavonols

**Supplementary figure 6. HPLC analysis of phenolic compounds extracted from flower tissue from wild type and FvMYB46-deletion mutants**. Individual phenolic compounds were grouped into; flavonols, flavan-3-ols and derivatives of ellagic and hydroxycinnamic acids.
