## Supplementary material for "CRISPR-Cas9 mediated deletions of *FvMYB46* reduces fruit set and biosynthesis of flavonoids in *Fragaria vesca*"

### Supporting methods

#### Identification and testing *F. vesca* U6-promoter sequences for gRNA expression

Using the AtU6-26-gene (X52528), the AtU6-1 gene (X52527) from *Arabidopsis*, and the *T. aestivum* U6 snRNA gene (X63066) in BLAST searches against the nr *Fragaria vesca* nucleotide database at NCBI or the Blast service at the Genome Database Rosaceae (GDR) against the *F. vesca* Whole Genome v4.0.a1 Assembly & Annotation, identified 8 unique U6 snRNA genes with promoters containing TATA boxes and USE (Upstream sequence elements) known to be important for expression (Supplementary figure 1). ~350 bp upstream sequences from the annotated TSS of these sequences comprising the putative U6-promoters were aligned with MEGA-6. The 329 bp upstream sequence of U6-1, 335 bp of the U6-2 upstream sequence and 335 bp of the U6-3 (acc nr) upstream sequence were selected for transient expression analysis in *F. vesca* berries to identify active promoters. The promoter sequences containing FvU6-BbsI-BbsI-gRNA-scaffold flanked by PacI, attR1 and EcoRI sequences were synthesized at GenArt (Thermo Fisher). https://www.thermofisher.com/no/en/home/life-science/cloning/gene-synthesis/geneart-gene-synthesis.html.The gRNA oligos for MET1 and RPPL1, were annealed to create 20 bp double stranded gRNAs with BsaI overhangs, and subsequently ligated into the BsaI cassette of the pMK-RQ-gRNA expression vectors linearized with BsaI. The U6-promoter-gRNA expression cassettes were then subcloned into the EcoRI site of pFGC - pcoCas9 vector for transient expression analysis.

*Transient expression analysis*

The constructs were transiently expressed in ripe *F. vesca* fruits by Agroinfiltration. qPCR on cDNA from RNA isolated from fruits 6 days after infiltration using gRNA specific and scaffold specific primers, showed that the FvU6-1 promoter showed the highest and most stable expression compared to the other *F. vesca* promoters and the AtU6-26 promoter suggesting this promoter would be the best option for stable expression of gRNA in *F. vesca*.

#### Screening putative CRISPR-Cas9-mutations and outcrossing of T-DNA

*Fragaria vesca* H4 leaf discs were transformed with pCAS9-TPC-FvU6-1-MYB46-2XgRNA using *Agrobacterium* mediated transformation. Developing calluses were propagated on MS-media containing 3 mg/L BASTA, and after shoots appeared, leafs were collected for PCR-screening using primers flanking the gRNA target regions to identify plants with deletions in *FvMYB46*. PCR screening identified 50 (?) plants with a deletion in the expected region suggesting successful deletion of the expected target. Of 50 primary transformants positive for BAR, 22 daughter plants (V1), each from different primary transformants, were analysed for genotyping. For V1 lines tested (n = 19) by PCR-genotyping with FvMYB46seqF and FvMYB46seqR2, a deletion in *FvMYB46* was inherited by all daughter plants positive for *CAS9* while no deletion was identified in *CAS9*-negative plants, suggesting that the deletion was inherited by the mother plants and coupled to the *CAS9*-gene (Figure 2 a). 3 primary transformants from the T0 generation (ID #43, #44 and #45) were selfed, and 24 T1 progeny plants were genotyped for deletions in *FvMYB46* and presence of *CAS9* using FvMYB46seqF and FvMYB46seqR2 and *CAS9*-primers (Cas9-F3n and Cas9-R3n) respectively. 23 of the plants gave a PCR-product of ~160bp suggesting that these plants carried the same deletions in *FvMYB46* as their respective parental line (Figure 2b, c). Surprisingly, plant #43_T1_16 gave a product of ~240bp, suggesting that it did not contain a mutation, suggesting that the mother plant originally was heterozygous for the deletion. 12 of the 23 T1 plants with a deletion in myb46 were negative for *CAS9*, suggesting successful outcrossing of the T-DNA.

As expected, the 43_T1_16 plant, which exhibited a non-deletion in *FvMYB46* did not show presence of *CAS9*

*Supplementary table 5. Oligos used in this study*

| Primer | Sequence (5´- 3´) |
|---|---|
| LB_Primary_1_R | TAATTGCGTTGCGCTCACTG |
| Pea3A_T_1_F | CCTCTCTTTGGTTATGTCTTGAATTGG |
| Cas9-F3n | CAACAACTACCACCACGCTCA |
| FvU6-26_2_F | CCTCTCAGGCCCAAACAGTC |
| LB_Nested_2_R | GAGAGGCGGTTTGCGTATTG |
| Cas9RTF | TGGTTTCGATTCTCCTACCG |
| Cas9-R3n | ATCCCTTCCCTTATCCCACAC |
| HindIII_flank_F_2 | GTCCGATTGGAAGCAAGAAC |
| Flank_upstream_T-DNA-2_F | AATGGTCAAAATACCACATAGGC |
| gRNA1_MYB46_OT1_F | CGTGTTCTTCCATCCTCACC |
| gRNA1_MYB46_OT1_R | ACACCGTCAGACTTCAATGG |
| gRNA1_MYB46_OT2_F | AACAACGGGGAAATAGAAGAGAG |
| gRNA1_MYB46_OT2_R | TGTAATGCAGGCACTTCCAC |
| gRNA1_MYB46_OT3_F | TTGCATGCATGTTGATGTTG |
| gRNA1_MYB46_OT3_R | TTCATGGTTCGCATTCACAG |
| gRNA2_MYB46_OT1_F | CTCCGGAAACCCAATAAGTG |
| gRNA2_MYB46_OT1_R | ACGAGGGTTTGTGTTTCGAG |
| FvMYB46seqF | GAACCCTATGCTGTAAGTAATACCG |
| FvMYB46seqR2 | ACCTCTCTTAAGGTCAGGTCTCAA |
| TPC_F | TCTTGAATTGGTTTGTTTCTTCAC |
| TPC_R | TAGACAAGCGTGTCGTGCTC |
| FvEF1ARTF | GCCCATGGTTGTTGAAACTTT |
| FvEF1ARTR | GGCGCATGTCCCTCACA |
| MYB46_qPCR_F | GAGAGGTGCGGAAAGAGTTG |
| MYB46_qPCR_R | CTGCAATTTGAGACCACCTG |
| sgRNA_F1 | TGTTTTAGAGCTAGAAATAGCAAGT |
| gRNA-Ra2 | GCACCGACTCGGTGCCAC |
